# Highly pathogenic avian influenza H5N1 virus infections in wild red foxes (Vulpes vulpes) show neurotropism and adaptive virus mutations

**DOI:** 10.1101/2022.07.21.501071

**Authors:** Luca Bordes, Sandra Vreman, Rene Heutink, Marit Roose, Sandra Venema, Sylvia B E Pritz-Verschuren, Jolianne M Rijks, José L Gonzales, Evelien A Germeraad, Marc Engelsma, Nancy Beerens

**Affiliations:** Wageningen Bioveterinary Research, Lelystad, the Netherlands; Dutch Wildlife Health Centre, Utrecht University, Utrecht, the Netherlands

**Author notes:** Address correspondence to Nancy Beerens.

## Abstract

During the 2020-2022 epizootic of highly pathogenic avian influenza virus (HPAI) several infections of mammalian species were reported in Europe. In the Netherlands, HPAI H5N1 virus infections were detected in three wild red foxes (*Vulpes vulpes*) that were submitted with neurological symptoms between December 2021 and February 2022. Histopathological analysis demonstrated the virus was mainly present in the brain, with limited or no detection in the respiratory tract and other organs. Phylogenetic analysis showed the three fox viruses were not closely related, but were related to HPAI H5N1 clade 2.3.4.4b viruses found in wild birds. In addition, limited virus shedding was detected suggesting the virus was not transmitted between the foxes. Genetic analysis demonstrated the presence of mammalian adaptation E627K in the polymerase basic two (PB2) protein of the two fox viruses. In both foxes the avian (PB2-627E) and the mammalian (PB2-627K) variant were present as a mixture in the virus population, which suggests the mutation emerged in these specific animals. The two variant viruses were isolated and virus replication and passaging experiments were performed. These experiments showed mutation PB2-627K increases replication of the virus in mammalian cell lines compared to the chicken cell line, and at the lower temperatures of the mammalian upper respiratory tract. This study showed the HPAI H5N1 virus is capable of adaptation to mammals, however more adaptive mutations are required to allow efficient transmission between mammals. Therefore, surveillance in mammals should be expanded to closely monitor the emergence of zoonotic mutations for pandemic preparedness.

**IMPORTANCE:** Highly pathogenic avian influenza (HPAI) viruses caused high mortality amongst wild birds in 2021-2022 in the Netherlands. Recently three wild foxes were found to be infected with HPAI H5N1 viruses, likely by feeding on infected birds. Although HPAI is a respiratory virus, in these foxes the viruses were mostly detected in the brain. Two viruses isolated from the foxes contained a mutation that is associated with adaptation to mammals. We show the mutant virus replicates better in mammalian cells than in avian cells, and at the lower body temperature of mammals. More mutations are required before viruses can transmit between mammals, or can be transmitted to humans. However, the infections in mammalian species should be closely monitored to swiftly detect mutations that may increase the zoonotic potential of the HPAI H5N1 viruses as these may threaten public health.

## INTRODUCTION

Since the introduction of highly pathogenic avian influenza virus (HPAI) H5 clade 2.3.4.4b in 2016, this virus clade caused repeated outbreaks in wild birds and poultry in Europe. Where low pathogenic avian influenza viruses replicate mostly in the digestive and respiratory epithelium with often mild disease in poultry and wild birds, these HPAI viruses can cause severe systemic disease including viremia leading to diffuse infection of several internal organs. Gallinaceous species are especially vulnerable to HPAI infection and high mortality is observed rapidly after infection. Infections with HPAI viruses of subtypes H5 and H7 are a notifiable disease in poultry and after detection of this virus a poultry flock is generally culled to prevent further spread. The 2020-2021 epizootic was a devastating outbreak for poultry and was followed by the 2021-2022 epizootic, which has become the largest HPAI outbreak in number of culled animals that ever occurred in Europe. In addition, high mortalities in an increasing number of species of wild birds was observed during both epizootics. Sporadically, HPAI H5 clade 2.3.4.4.b infections of free-living wild carnivore species were reported besides the infections in wild birds and poultry. Late 2020, a disease and mortality event involving four juvenile common seals (*Phoca vitulina*), one juvenile grey seal (*Halichoerus grypus*) and a red fox (*Vulpes vulpes*) at a wildlife rehabilitation center in the United Kingdom was reported to be associated with HPAI H5N8 infection (1). In May 2021, HPAI H5N1 infection was reported in two cubs of red foxes in the Netherlands (2). In August 2021, HPAI H5N8 infection was detected in three adult harbor seals (*Phoca vitulina*) found at the German North Sea coast (3). Interestingly, neurological signs were reported for several of these mammals and virus was detected in brain tissue. For mammals the most probable route of infection is ingestion of contaminated water, feces or infected bird carcasses. Foxes were infected experimentally via consumption of chicken carcasses infected with clade 2.2 HPAI H5N1, which underlines the possibility to infect wild mammals with HPAI via infected bird carcasses (4). Similarly, experimentally infected cats were also susceptible to HPAI H5N1 (A/Vietnam/1194/2004) via infected bird carcasses. Both the gut and myenteric plexus (5) and the olfactory bulb (6) were suggested as potential sites of virus entry into the central nervous system after consumption of infected birds.

The species barrier between birds and mammals is considerable. Therefore, adaptation of HPAI virus is needed for efficient replication and transmission in mammals. Current studies suggest at least three requirements for (efficient) airborne transmission of HPAI viruses between mammals: (i) efficient attachment of the viral hemagglutinin (HA) glycoprotein to α2,6-linked sialic acid receptors, which are present in the upper respiratory tract of mammals. Mutations in HA are required to change the HA binding properties from the avian-type α2,3-linked sialic acid receptors to the mammalian-type receptor (7). (ii) Optimal stability of HA protein in mammalian airways. Mutations in HA are required to optimize fusion of the viral and endosomal membranes and the subsequent release of the viral genome in the cytoplasm (8). (iii) Increased virus replication through mammalian adaptation substitutions in the polymerase complex (9, 10). Although these adaptations favored airborne transmission of the studied HPAI H5N1 strain A/Vietnam/1204/2004, it remains to be elucidated whether these findings can be extrapolated to other clades of HPAI H5N1 viruses, such as the current HPAI H5N1 clade 2.3.4.4b virus. However, mutation E627K has also been detected in the polymerase basic protein 2 (PB2) of HPAI H5 clade 2.3.4.4b viruses isolated from mammals in 2021. Two out of three seals infected with HPAI H5N8, detected in August 2021 in Germany, contained the PB2-E627K mutation (3). The PB2-627K variant has been identified as an adaptation of the virus polymerase machinery, that likely stimulates virus replication at lower temperatures of the upper respiratory tract in mammals (9, 10). For HPAI H5 viruses enhanced virus replication caused by PB2-E627K has been shown to increase pathogenicity *in vitro* and *in vivo* in mice (11-13). Furthermore, the recently discovered mammalian infections of the fox and seals in the UK late 2020 contained the PB2-D701N adaptation, which was also suggested to increase virus replication (9, 14). These findings may suggest the current HPAI H5 clade 2.3.4.4b viruses have an increased zoonotic potential and are able to infect mammals resulting in adaptations in the PB2 gene segment.

The previously described cases of HPAI H5 clade 2.3.4.4b infections in seals and foxes and the three infected foxes detected in the Netherlands between December 2021 and February 2022 suggest the incidence of infections in mammals might be increasing. In this study, we analyzed virus localization in tissues from the three foxes with related histopathology and showed the virus is mainly present in the brain, with limited detection in the respiratory tract. Phylogenetic analysis showed the three fox viruses were not closely related, but were related to viruses found in wild birds. Genetic analysis demonstrated the presence of mutation PB2-E627K in two out of three fox viruses. From one of these foxes two virus variants were isolated, one containing the avian PB2-627E variant and one containing the mammalian PB2-627K variant. Virus replication and stability of both isolates was studied over time and showed increased replication for the PB2-627K variant in human and dog cell lines compared to the chicken cell line and at the temperature of the mammalian upper respiratory tract compared to the temperature of the avian respiratory tract, indicating more efficient replication *in vitro* in mammals compared to birds.

## RESULTS

### Virological analysis of infected foxes

Three foxes were found in the Netherlands at the municipality Dorst, Heemskerk and Oosterbeek showing abnormal behavior. Clinical signs in fox-Dorst were apparent blindness, head shaking, falling over and opisthotonos. This animal was euthanized on 3/12/2021. Fox-Heemskerk showed convulsions, as reported, and was euthanized on 1/1/2022. Fox-Oosterbeek showed lethargy, crouched down with convex back and legs tucked under. The animal lacked fleeing behavior but was alert and did turn its head to look when approached. This fox was humanely dispatched on 7/2/2022. The distance between the different locations was 80km to 105km (Figure 1).

**Figure 1:**
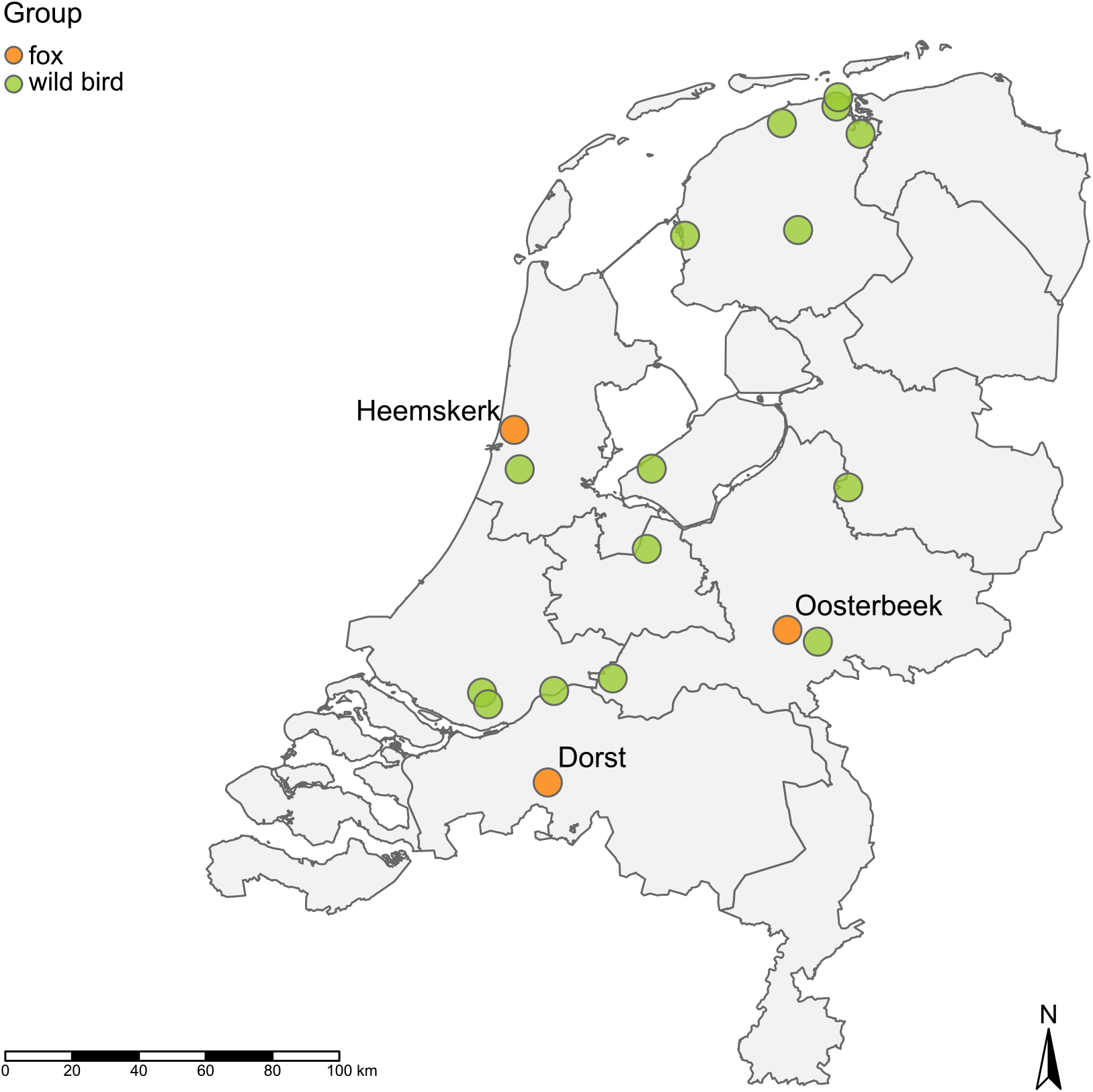
Locations of the three infected foxes and wild birds from which closely related viruses were isolated during the 2021-2022 HPAI H5N1 epizootic in the Netherlands.

The foxes were submitted for testing on avian influenza virus, tracheal and rectal swabs were taken and brain tissue was collected. Interestingly, the brain samples from two different locations (amnion horn and medulla oblongata, cerebrum and cerebellum) tested positive for avian influenza virus by M-PCR with high virus genome loads, whereas no virus was detected in rectal swabs, and considerably lower virus loads were detected in the throat swabs of two foxes. Only for fox-Heemskerk comparable virus loads were detected in the brain samples and throat swab (Table 1). The viruses were subtyped as HPAI H5N1 using Sanger sequencing. Virus isolation was performed successfully on brain tissue of all three foxes, showing the presence of infectious virus in the brain.

**Table 1:**
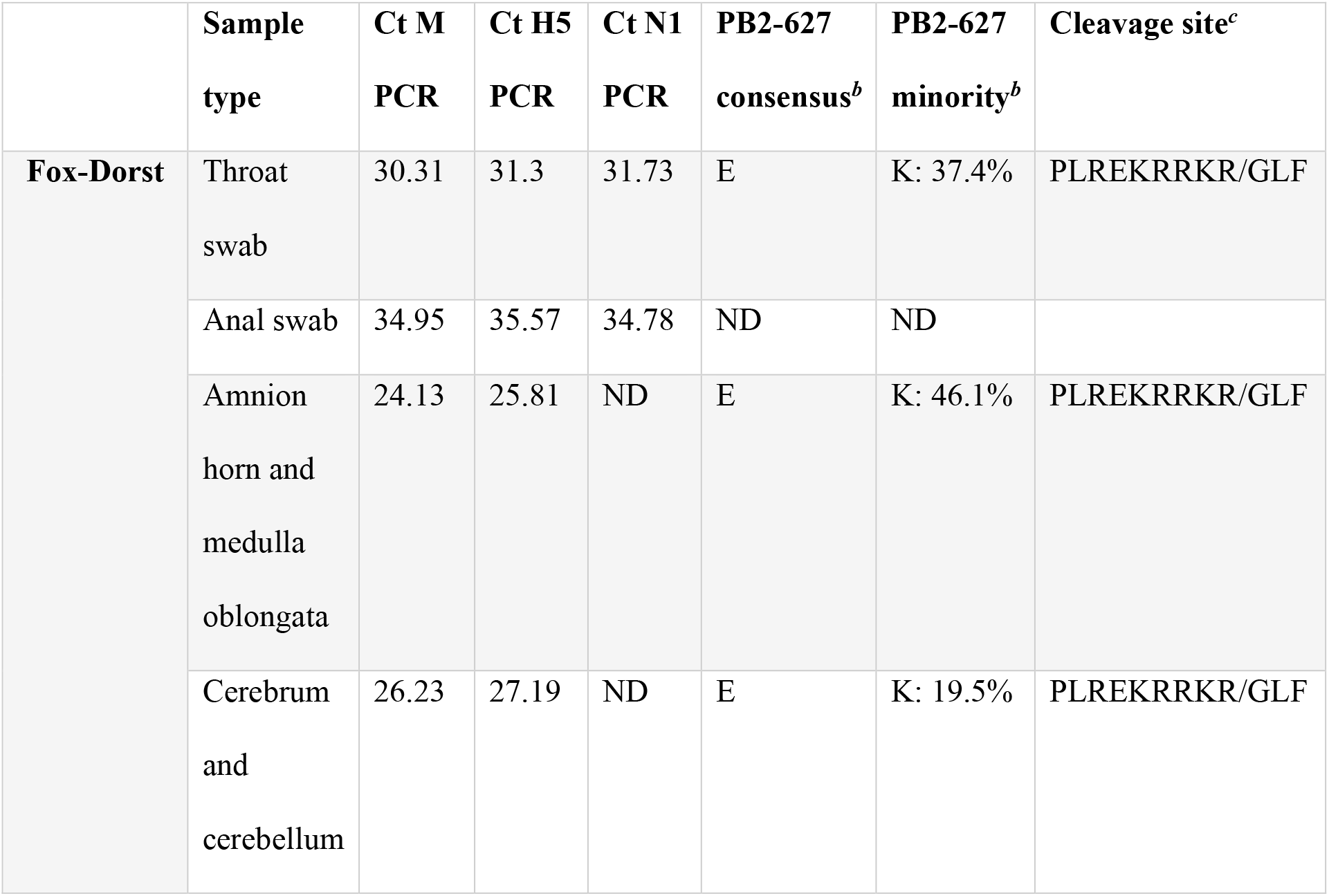
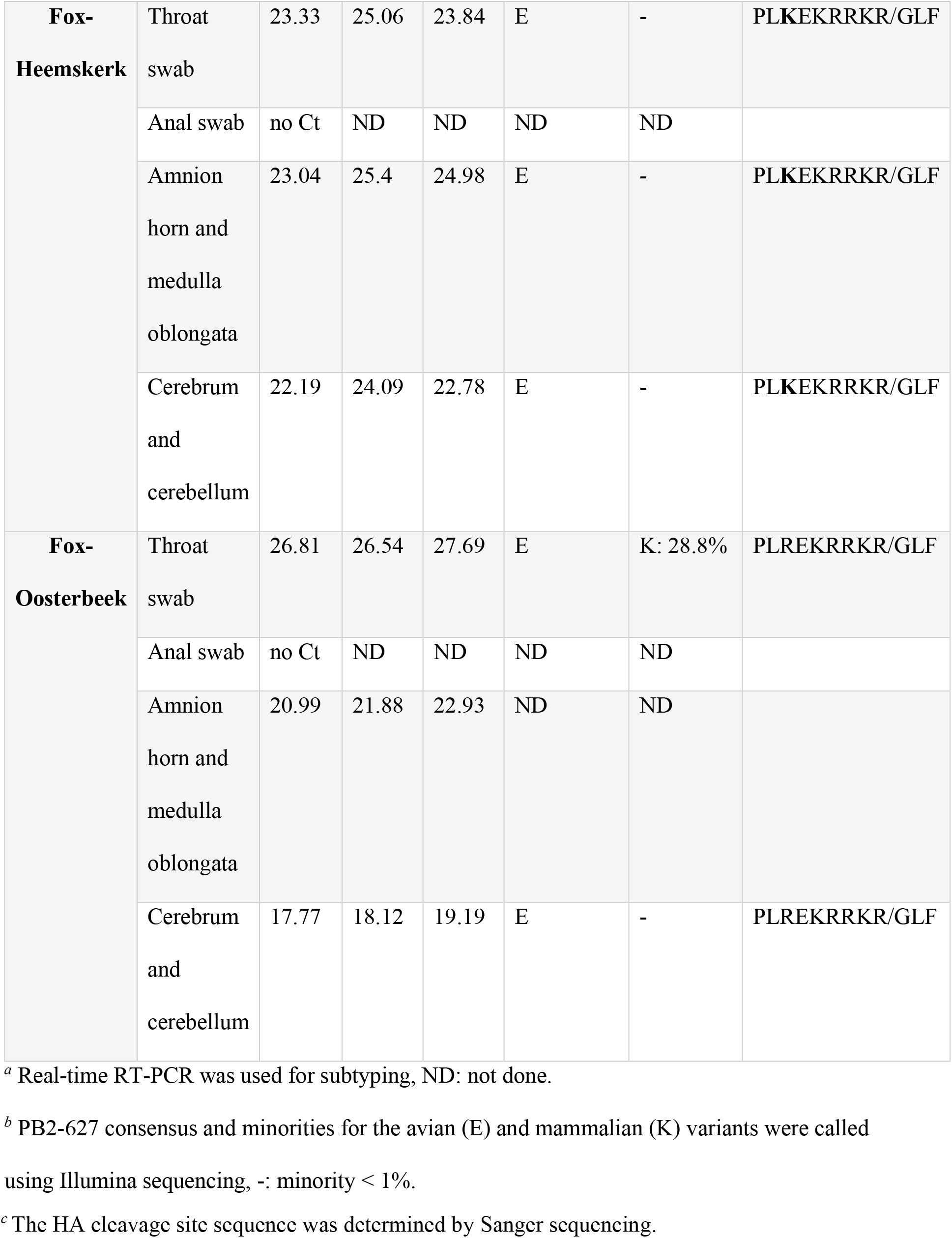
Virus detection, Hemagglutinin/Neuraminidase-subtyping and sequencing.^*a*^.

**Table 2:**
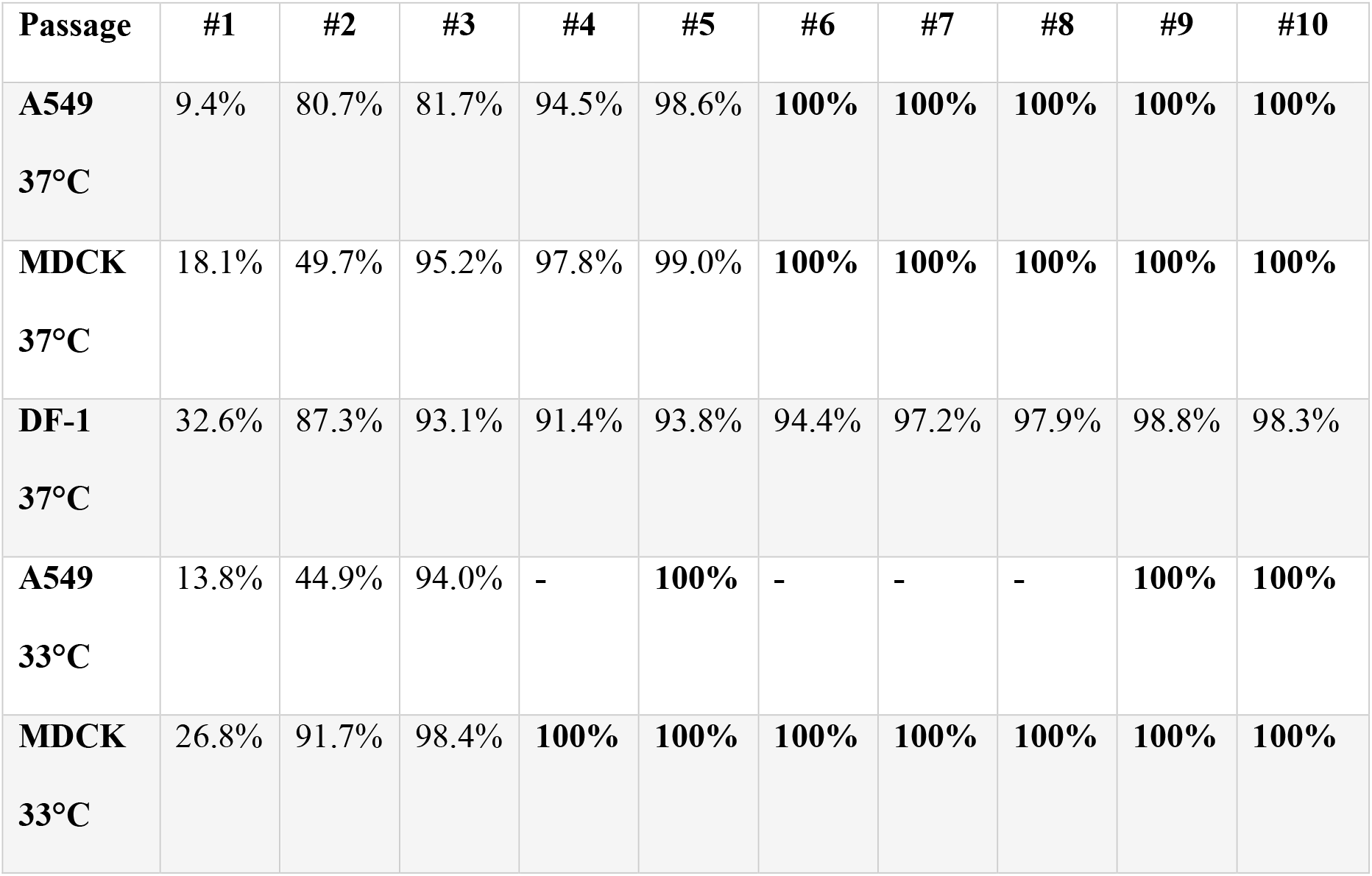
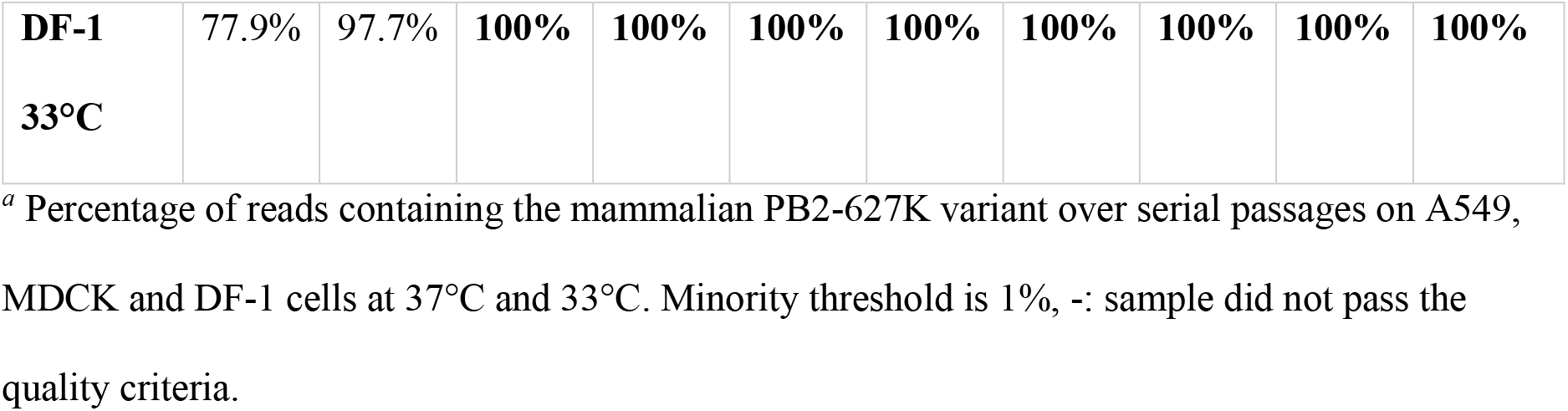
Serial passaging of the PB2-627 variants on cell lines.^*a*^.

### Pathological examination of infected foxes

All three foxes were adult males with a moderate to poor body condition. Two foxes had a shiny coat and one fox (Oosterbeek) was covered with mud. The most prominent gross finding were poorly collapsed lungs with a marbled red aspect, which was present with a slight variation in all three foxes. Histology revealed a subacute chronic purulent to granulomatous broncho-interstitial pneumonia with large numbers of parasitic structures (*Angiostrongylus vasorum*) in two foxes. These pulmonary changes were not associated with virus protein expression (Figure 2A and B and Table S1). The upper respiratory tract (trachea and nasal conchae) displayed a mild to moderate suppurative inflammation with presence of parasitic structures (*Capillaria spp*.). In fox-Heemskerk virus protein expression was observed in the olfactory epithelial cells of the nasal conchae (Figure 2C), while there was no protein expression in the trachea and nasal conchae of the other foxes (Figure 2D). In all three foxes strong virus protein expression was present in the brain, which was most prominent in the cerebrum (Figure 2E and F). Virus protein was expressed in neurons and microglia cells in the neuropil and was associated with non-suppurative polioencephalitis with perivascular cuffing. There was no virus expression or significant histopathology in the olfactory bulb (Figure 2G). A subacute lymphoplasmacytic myocarditis with myocardial degeneration and necrosis with virus protein expression in cardiomyocytes was detected only in fox-Heemskerk (Figure 2H). No virus protein expression was observed in the other investigated organs (intestinal tract (Figure 2I), pancreas, spleen, liver and kidney (Table S1)).

**Figure 2:**
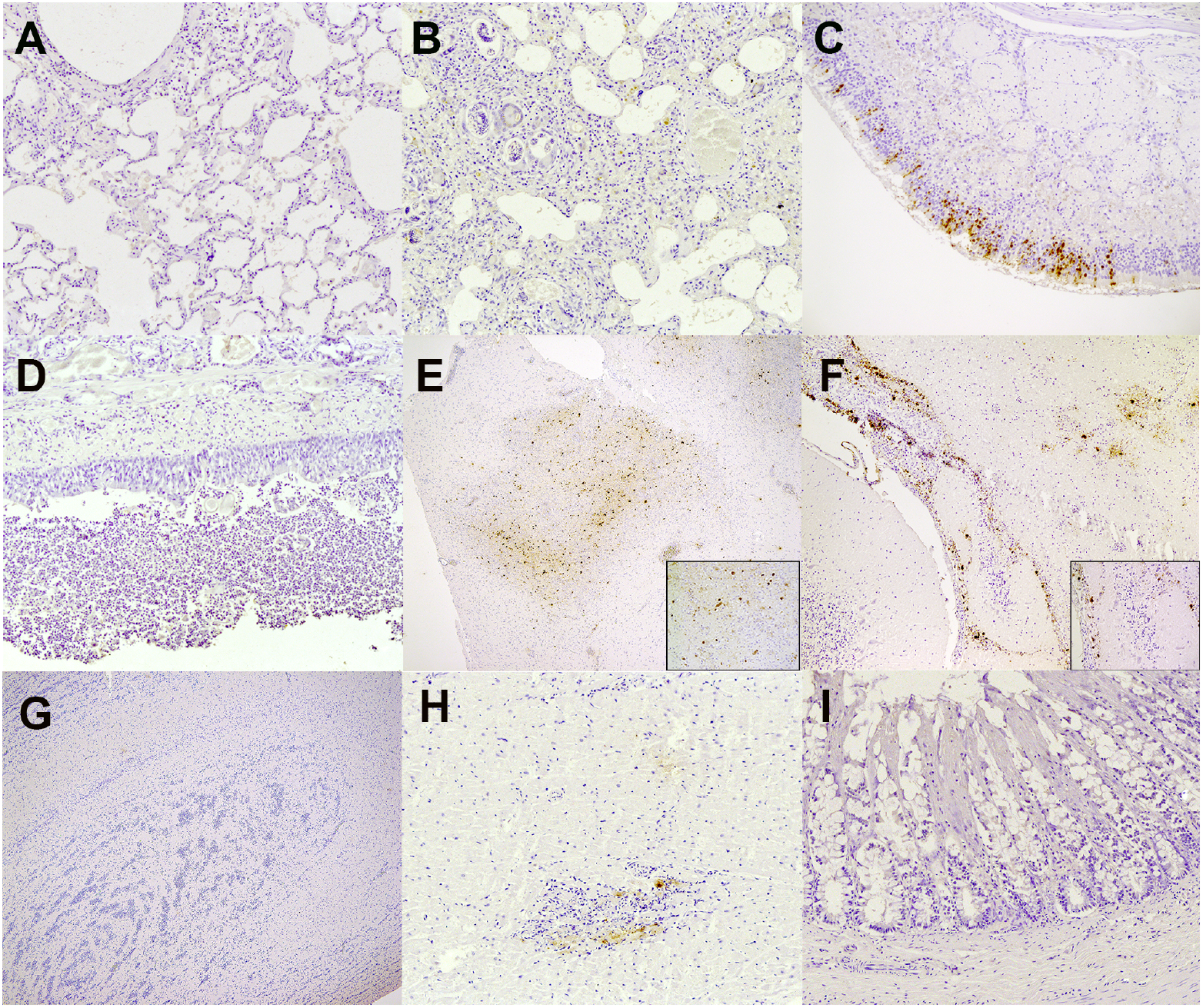
Histopathology and virus protein expression in tissues of foxes. Immunohistochemistry (IHC) was performed on tissue sections against influenza A nucleoprotein, insets show hematoxylin and eosin stain (HE) or magnification of IHC stain A) Fox-Dorst moderate purulent broncho-interstitial pneumonia (less affected area), no virus protein expression; B) Fox-Oosterbeek severe purulent broncho-interstitial pneumonia with intralesional larvae (*Angiostrongylus vasorum*), no virus protein expression; C) Fox-Heemskerk mild to moderate purulent rhinitis with positive staining of olfactory epithelial cells; D) Fox-Dorst severe necropurulent tracheitis with intralesional parasite eggs (*Capillaria spp*), no virus protein expression; E) Fox-Dorst cerebrum moderate non-suppurative polioencephalitis with virus protein expression in neurons and microglia cells in the neuropil; F) Fox-Heemskerk cerebellum moderate non-suppurative polioencephalitis with virus protein expression in neurons and microglia cells in the neuropil; G) Fox-Oosterbeek bulbus olfactorius no significant histopathology and virus protein expression; H) Fox-Heemskerk moderate lymphoplasmacytic myocarditis with mild myocardial degeneration and necrosis with positive virus protein staining of cardiomyocytes; I) Fox-Oosterbeek colon, no significant histopathology and virus protein expression; A, B, C, D, H and I magnification 20x, F-magnification 10x, E-G, original magnification 2,5x – insets, magnification 40x.

### Phylogenetic and genetic analysis of fox viruses

Full genome sequencing was performed on brain samples and throat swabs to study the genetic relationship between the viruses. Phylogenetic analysis showed that the viruses belonged to H5 clade 2.3.4.4b, and clustered with viruses found in wild birds during the HPAI H5N1 2021-2022 epizootic in the Netherlands. The viruses had the same genetic constitution as the HPAI H5N1 viruses found in the Netherlands during the 2020-2021 epizootic. Closest related wild bird viruses differed between 17 and 42 nucleotide positions from the fox viruses and were found between 39 and 149 km distance, 5 to 37 days before the infected foxes were detected (Table S2). The viruses isolated from the foxes were not closely related based on the phylogenetic analysis and differed between 127 and 176 nucleotides from each other (Figure 3 & Figure S1). Therefore, the three foxes were likely infected by independent introductions from wild birds.

**Figure 3:**
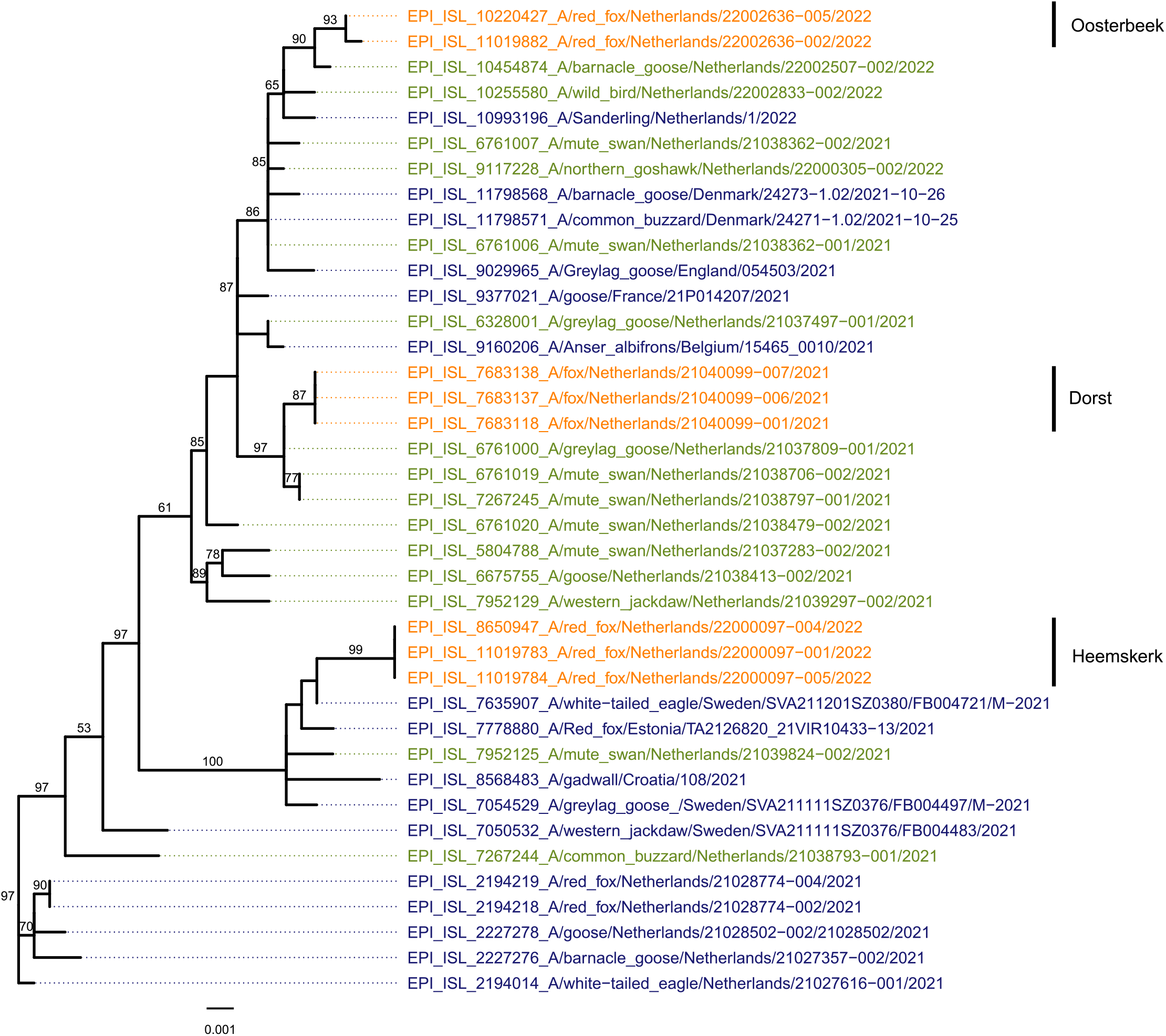
Phylogenetic tree for the HA segment obtained with the Maximum Likelihood method showing the viruses detected in the samples of the three foxes (orange), and closely related HA sequences from other viruses detected in the Netherlands (green), Europe (blue) and relevant sequences form the 2020-2021 epizootic (blue).

Genetic analysis was performed to investigate whether mutations implicated in adaptation to mammalian hosts occurred in the virus genomes. Mutation screening identified the previously described E627K mutation in the PB2 segment of the fox-Dorst virus. A minority variant analysis of the next-generation sequencing data from three samples of fox-Dorst showed a mixture of the avian (627E) and mammalian (627K) PB2 variant. The percentage of the 627K mutation varied between 37.4% of the virus population in the trachea swab, 46.1% in the amnion horn and medulla oblongata and 19.5% in the cerebrum and cerebellum (Table 1). Fox-Heemskerk carried only the avian 627E-variant, and fox-Oosterbeek carried the avian 627E-variant with 28.8% of the 627K-variant in the throat swab. Further in-depth analysis of the sequencing data did not reveal other known or previously described host shift adaptations in the fox viruses (results not shown).

### The role of mutation E627K in replication of the fox-Dorst virus

A limited dilution series of the amnion horn and medulla oblongata sample derived from fox-Dorst was inoculated into embryonated eggs for virus isolation. Full genome sequencing of the viruses was performed for the viruses isolated from the individual eggs. This showed we isolated a HPAI H5N1 virus with the mammalian PB2-627K variant (100%), and a virus with the avian PB2-627E variant (98,2%) containing only one additional mutation (G485R) in the nucleoprotein (NP) (99%). To study the effect of PB2-E627K on virus replication, the virus titre was measured at specific time points after infection of mammalian A549 and MDCK cell lines and the avian DF-1 cell line.

The mean virus titre appears to increase from 48h post infection (p.i.) onwards due to the PB2-E627K mutation on the mammalian cells (A549, MDCK) but not the avian cells (DF-1) at 37°C (Figure 4). The putative differences between the two virus variants were statistically assessed using fitted linear mixed models (LMM), with post hoc analysis. Due to sample size limitations, three way interactions containing time, virus and cell type as a variable could not be analyzed. Thus, differences between cell types could not be investigated using the LMM. However, the two way interactions of time with virus, and time with temperature, could be assessed using the LMM. The differences between the two virus variants were statistically significant from 48h p.i. onwards (p < 0.05)(Table S3). To assess the apparent absence of differences between the PB2-627E and PB2-627K virus replication curves on DF-1 cells at 37°C virus titre was compared by performing pair-wise comparisons between both viruses at each time point and ignoring the data dependency (no random effects introduced in the model) due to replicated measures. This pairwise comparisons showed no significant difference (p > 0.05) between the two viruses from 48h p.i. onwards. These results may indicate that PB2-E627K enhances virus replication in mammalian cells but not avian cells at the temperature of the avian upper respiratory tract (37°C).

**Figure 4:**
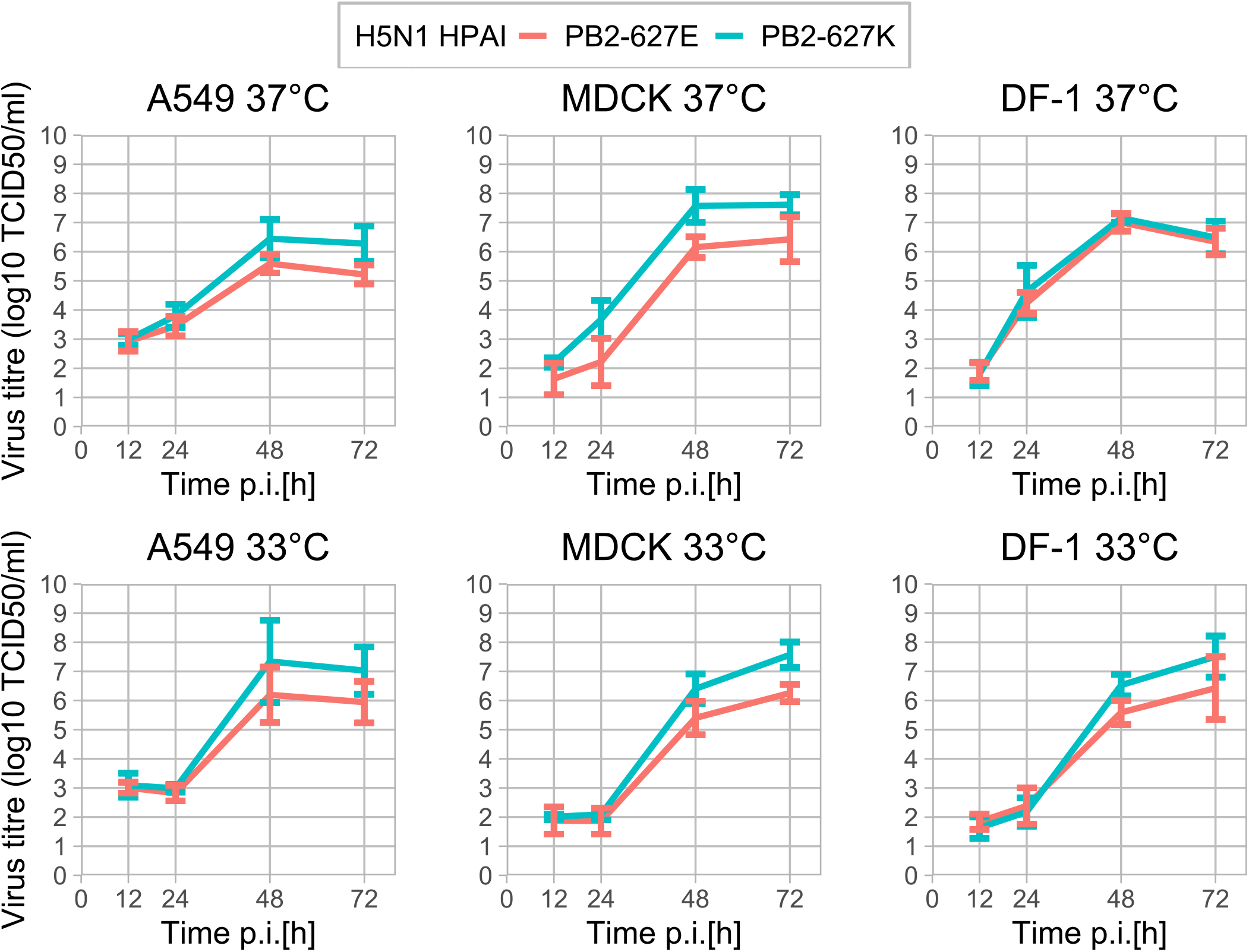
Virus replication curves of the PB2-627K (blue) and PB2-627E (red) H5N1 HPAI viruses on A549 (human), MDCK (dog) and DF-1 (chicken) cells cultured at 37°C, the temperature of the avian upper respiratory tract and 33°C, the temperature of the mammalian upper respiratory tract. Differences in infectious virus titre between viruses is significant from 48h p.i. onwards (p < 0.05). No significant differences were found between infectious virus titre of the two viruses on DF-1 cells at 37°C (p > 0.05). Virus titre on cells cultured at 33°C instead of 37°C were significantly lower at 24h p.i. (p < 0.0001) and 48h p.i. (p < 0.05) but significantly higher at 72h p.i. (p < 0.05).

The PB2-E627K mutation is likely an adaptation of the polymerase to the lower body temperature of mammals compared to birds. Therefore, virus replication was also studied at 33°C, which is the approximate temperature of the mammalian upper respiratory tract. The LMM analysis of the variables time and temperature indicated virus replication was significantly lower at 24h p.i. (p < 0.0001) and 48h p.i. (p < 0.05) but significantly higher at 72h p.i. (p < 0.05) indicating the replication of both virus variants is delayed but not impaired (Figure 4 and Table S4). In contrast with our findings at 37°C virus titre was significantly higher for the PB2-627K variant then the PB2-627E variant on DF-1 cells cultured at 33°C (Figure 4 and Table S3). Thus, PB2-E627K increases virus replication on mammalian cells and avian cells at the temperature of the upper respiratory tract of mammals (33°C).

### Serial passaging of the two virus variants

Serial passages on the mammalian A549 and MDCK cell lines and the avian DF-1 cell line cultured at 37°C and 33°C were performed to assess the stability of the obtained virus isolates. The 100% isolate of PB2-627K was stable on all experimental conditions for 10 passages in all cell lines at both temperatures tested (data not shown). However, passaging of the PB2-627K virus isolate, which contains 1.8% PB2-627K showed the 627K mutant has a replication benefit in all three cell lines at both temperatures tested. However, whereas the 627K-mutation is selected in the mammalian cell lines within six passages at 37°C, selection in the avian DF-1 cell line takes more than 10 passages. These results suggest the PB2-E627K mutation increases replication capacity of the virus in mammalian cells compared to avian cells. Furthermore, PB2-627K was selected faster (within 3 or 4 passages) on cells incubated at 33°C compared to cells incubated at 37°C. Thus, the lower temperatures of the mammalian upper respiratory tract (33°C) could be involved in the selection of the PB2-E627K mutation.

## DISCUSSION

Here we investigated three cases of HPAI H5N1 infections in wild foxes in the Netherlands to assess tissue tropism in wild mammals and screen for adaptive mutations. All three foxes showed unusual tissue tropism and evidence was found for mammalian adaptation. Full genome sequencing of the fox viruses followed by phylogenetic analysis demonstrate they belong to clade 2.3.4.4b and are related to viruses detected in wild birds. The fact that the three fox-viruses were not closely related, and the large distance between the locations at which the three foxes were found suggest these were separate virus introductions likely originating from wild birds. Experimental infection by consumption of infected bird carcasses has indicated foxes are susceptible to AIV infections, which is also the most probable route of infection for free-living foxes (4). Virus RNA was most abundant in the brain of all three foxes and was associated with positive IHC staining for virus protein in brain tissue of all three foxes. This finding is novel, as previous HPAI H5 virus infections in foxes showed a diffuse tropism with high virus replication in the respiratory system similar to HPAI infection in poultry (1, 4). However, it is unclear how the virus infected the brain without clear viremia and systemic replication. Fox-Heemskerk showed mild positive staining by IHC of the olfactory epithelium of the nasal conchae. We also detected high viral RNA loads in the throat swab of this fox most likely originating from the olfactory epithelium since all other respiratory organs were tested negative by IHC. The olfactory epithelium is connected to the brain via the olfactory bulb and has been previously described as a point of entry for AIV (5, 6). However, we found no evidence of AIV replication in the olfactory bulb based on IHC and absence of histopathologic changes, while in other parts of the brain, virus expression was always associated with histopathologic changes. The olfactory epithelium could play a role in virus entry but the exact route of infection remains to be elucidated. Fox-Heemskerk also showed mild positive virus staining of cardiomyocytes. Infection of cardiomyocytes normally indicates a systemic disease and viremia. Currently, we have no information on the early stages of infection, or infections that may occur without display of neurological symptoms in wild foxes. Thus, we cannot exclude virus replication also occurred in the respiratory tract or other organs at a particular stage of the infection. However, in these foxes showing neurological symptoms exclusive neurotropism of the virus was observed. Neurological symptoms have also been reported for avian species. For example, HPAI disease in chickens is short and results in sudden death however, in domestic ducks and wild birds disease is longer and often neurological symptoms like partial paralysis and tremors can be observed (15-17). However, HPAI is generally considered to be a respiratory disease with high virus genomic RNA concentration in the respiratory system (18). Thus, the lack of virus replication in the respiratory system of these foxes is interesting and suggests the current HPAI H5N1 2021-2022 viruses have an increased neurotropism in mammals.

In two of the three foxes, a minority population of viruses was identified containing the zoonotic mutation PB2-E627K. The fact that this mutation was not detected in any of the wild bird sequences during the 2021-2022 epizootic in the Netherlands suggests this mutation quickly arises upon infection of mammals. Furthermore, as both the PB2-627E and PB2-627K variants were detected in these two foxes, it appears likely that this mammalian adaptation emerged within these specific animals. A similar event has occurred in two seals infected with HPAI H5N8 virus in August 2021 in Germany, and indicates selection pressure for this adaptation in mammals (3). Histopathology revealed distinct virus protein expression and associated brain histopathology with no virus replication found by IHC in the lungs in these seals, similar to the foxes that were investigated in this study. We also showed mutation PB2-E627K increased replication in the mammalian cell lines at both 33°C and 37°C, whereas for the avian DF-1 cell line this was not observed at 37°C. Furthermore, in passaging experiments mutation PB2-627K was found to have a strong replication benefit in the tested mammalian cell lines at 37°C, whereas this effect was smaller on the avian cell line at this temperature. At 33°C the mutation PB2-627K was found to increase the replication capacity of the virus in all three cell lines to similar levels. Thus, mutation PB2-E627K increases replication speed on mammalian cells but not avian cells at relevant temperatures of the upper respiratory tract (33°C). This finding is supported by previous reports on the PB2-E627K mutation (9, 10). Therefore, the lower temperatures of the mammalian upper respiratory tract could be a driving factor for emergence of the PB2-E627K mutation. Although PB2-E627K improves virus replication in mammalian cells, the mutation appears not essential for virus replication on mammalian cells, which is in agreement with the initial introduction of the HPAI H5N1 virus carrying the PB2-627E variant from wild birds. The additional mutation G485R in the NP that is present in the isolated PB2-627E virus could be associated with adaptation to mammals as described in a previous study (19), however its effect on virus replication is not known. Although a relatively small increase in replication capacity was measured on mammalian cells due to the PB2-E627K mutation, previous studies have indicated small increases in HPAI virus replication coincided with increased pathogenicity in mammals (11, 12). Increased virus replication may also stimulate the emergence of further mammalian adaptations as more genomic copies are produced per introduction.

It is currently unclear which factors have contributed to the increase of infections observed in red foxes in nature. The HPAI H5N1 clade 2.3.4.4b virus may be more capable of infecting mammals, the virus may be more infectious or there may be a higher prevalence of the virus in wild birds during the 2021-2022 epizootic compared to previous epizootics. Alternatively, the increased neurotropism of the virus may have contributed to the infection of wild red foxes. Limited virus shedding and virus replication was observed in the respiratory system and digestive tract of the investigated foxes, which likely limits transmission between mammals. Consistent with this, no evidence for transmission between foxes was found based on the phylogenetic analysis of the viruses. Genetic analysis suggests that the zoonotic mutation PB2-E627K may arise in infected mammals. Although PB2-E627K is an important mammalian adaption, previous research has indicated several further adaptations are required for efficient air-borne transmission of AIV between mammals. For example, the PB2-E627K, HA-Q222L and HA-G224S mammalian adaptations were required before ten serial passages in ferrets to eventually produce airborne variants of HPAI (20). In particular adaptations in the viral HA glycoprotein, which affects the stability in the mammalian airways and receptor binding specificity to the α2,6-linked sialic acid receptors in the mammalian upper respiratory tract, should be monitored closely to prevent potential spread between mammals. Unfortunately, infections of mammalian carnivorous wild-life are difficult to prevent during HPAI epizootics when large numbers of wild birds are affected. Clearing of wild bird carcasses could help limit these HPAI introductions into wild mammals, but may not be feasible during large outbreaks and at remote sites in nature. Awareness should be raised for the potential transmission of HPAI viruses from wild birds to pet animals. Dogs and cats may be at risk when interacting with or feeding on infected wild birds or their carcasses. The observed tissue tropism in these foxes and the lack of evidence for further spreading between wild mammals indicates that it is unlikely HPAI H5N1 clade 2.3.4.4b spreads to humans. However, surveillance for HPAI viruses in wild mammals should be expanded to closely monitor the emergence of zoonotic mutations for pandemic preparedness.

## MATERIALS AND METHODS

### Tissue sampling, virus detection and histopathology

Three foxes showing neurologic signs were submitted for necropsy to exclude rabies and influenza A virus infection according to governmental surveillance guidelines. A post-mortem examination was performed within two days after euthanasia. Various tissue samples were taken for histopathology and immunohistochemistry (IHC) and fixed in 10% neutral buffered formalin. Tissues were processed and evaluated for histopathologic changes with haematoxylin and eosin stain (HE) and for influenza A nucleoprotein expression with IHC as described previously (15). From all animals an anal swab, throat swab and brain sections (amnion horn and medulla oblongata, cerebrum and cerebellum) were collected for viral RNA isolation. Swabs collected during post-mortem examination were placed in 2 ml of Tryptose Phosphate Broth (TFB) supplemented with 2.95% gentamycin. Avian influenza virus (AIV) RNA was subsequently extracted using the MagNA Pure 96 system (Roche, Basel, Switzerland). AIV was detected by a quantitative real-time RT-PCR targeting the matrix gene (M-PCR), as described previously (21). Positive samples were subtyped using H5-and N1-specific real time RT-PCRs as recommended by the European Union reference laboratory (22, 23). For at least one sample of each fox the HA cleavage site sequence and the N subtype were determined by Sanger sequencing as described elsewhere (21).

### Complete genome sequencing and analysis

All virus genome sequences were determined directly on the swab or tissue samples. Virus RNA was purified using the High Pure Viral RNA kit (Roche, Basel, Switzerland), amplified using universal eight-segment primers and directly sequenced, as described previously (21). Purified amplicons were sequenced at high coverage (average > 1000 per nucleotide position) using the Illumina DNA Prep method and Illumina MiSeq 150PE sequencing. The reads were mapped using the ViralProfiler-Workflow, an extension of the CLC Genomics Workbench (Qiagen, Germany). Consensus sequences were generated by a reference-based method. Reads were first mapped to a reference set of genomes, and subsequently remapped to the closest reference sequence. Finally, the consensus sequence of the complete virus genome was extracted and minority variants were called using a cutoff of 1%.

### Phylogenetic analysis

In addition to the virus sequences obtained from foxes and wild birds in the Netherlands in this study, the top-5 BLAST results for related sequences in Eurasia were included in the phylogenetic analysis. These H5N1 genome sequences were downloaded from the GISAID database (24). Phylogenetic analysis of the complete genome sequences was performed for each genome segment separately, aligning the virus sequences using MAFFT v7.475 (25), reconstructing the phylogeny using maximum likelihood (ML) analysis with IQ-TREE software v2.0.3 and 1000 bootstrap replicates (26). ML tree was visualized using the R package ggtree (27). The GISAID sequences used in the phylogenetic analysis are listed in Table S5, in which we acknowledge all contributors to the GISAID database.

### Cell cultures

Madin-Darby Canine Kidney (MDCK) cells obtained from Philips-Duphar (Weesp, the Netherlands), chicken embryo fibroblasts (DF-1)(ATCC, Wesel, Germany) and human lung alveolar epithelial cells (A549)(ATCC, Wesel, Germany) were maintained in cell culture medium consisting of Dulbecco’s Modified Eagle Medium GlutaMAX (DMEM)(Thermo Fisher Scientific, Bleiswijk, Netherlands) supplemented with 5% fetal calf serum (Capricorn scientific, Germany) and 0.1% penicillin-streptomycin (Thermo Fisher Scientific, Bleiswijk, Netherlands) at 37°C, 5% CO_2_. Cells were passaged when confluent using 0.05% Trypsin-EDTA (Thermo Fischer Scientific, Bleiswijk, Netherlands).

### Virus isolation, titration and propagation

Tissue homogenates from brain tissue (Amnion horn and medulla oblongata, cerebrum and cerebellum) were incubated for 1 hour with 1% penicillin and 1% gentamycin at room temperature. The tissue homogenates were filtered and injected in 9-day-old specific pathogen free (SPF) embryonated chicken eggs (ECE), as described previously (28). Allantoic fluid was harvested from deceased eggs, aliquoted, stored at -80°C and sequenced using Illumina sequencing as described above. Subsequently, a serial dilution of the amnion horn and medulla oblongata homogenate was injected in fresh 9-day-old SPF ECE’s. Allantoic fluid was harvested from all eggs and individually sequenced. The median tissue culture infective dose (TCID50) of the isolated viruses was determined by end-point titration on MDCK cells. In short, 2.5×10^4 cells per well of a 96-well plate were seeded overnight in cell culture medium. Monolayers were infected with ten-fold serial dilutions in infection medium consisting of DMEM glutaMAX supplemented with 0.1% penicillin-streptomycin and 0.3% bovine serum albumin. Each dilution was tested in eight-fold and each titration was diluted in triplicate. After two days of incubation at 37°C, 5% CO_2_ the monolayers were stained by immunoperoxidase monolayer assay (IPMA). Monolayers were fixed with 10% neutral-buffered formalin, primary antibody was produced in-house from HB 65 mouse anti-nucleoprotein and diluted 1:2500 followed by HRP-conjugated rabbit anti-mouse diluted 1:500 (Dako, Glostrup, Denmark). The titration was repeated on a different day and TCID50 titres were calculated using Reed and Muench (29).

### Virus replication

The multiplicity of infection (MOI) was optimized for each cell line by selecting a dilution, which caused limited cell death but sufficient virus replication. MDCK, DF-1 and A549 cells were seeded in duplo on a 24-well plate at a density of 2.5×10^5 cells per well in cell culture medium. The next day, cells were infected with 1.5 ml of virus diluted in infection medium at a multiplicity of infection of 0.01 (A549), 0.001(MDCK) or 0.0005(DF-1) and incubated at 37°C, 5% CO_2_ or 33°C, 5% CO_2_. At indicated time points, 150 μl of medium was harvested and stored at -80°C. 150 μl of fresh infection medium was added after each harvest. Infectious virus titre of the collected medium was determined in triplo using the TCID50 protocol described above.

### Passaging of viruses and sequence analysis

Stability of the virus isolates was determined by passaging ten times on A549, MDCK and DF-1 cells at identical temperatures and MOI as the replication experiment described above. In short, cells were seeded at a density of 2.5×10^6 cells per T-25 flask. The next day, cells were infected with diluted virus in 3 ml infection medium. Cells were incubated either at 37°C, 5% CO_2_ or 33°C, 5% CO_2_ for three days before the medium was collected. The medium was diluted 1000x and added to a fresh T-25 flask of A549, MDCK or DF-1 cells. The remaining undiluted medium was stored at -80°C for sequencing. Viral RNA was isolated with the Zymo Quick-RNA Viral 96 Kit (BaseClear, Leiden, Netherlands). A region of 168 base pairs on the PB2 protein was amplified by RT-PCR using custom primers (Table S6)(Eurogentec, Maastricht, Netherlands) and the QIAGEN OneStep RT-PCR Kit (Qiagen, Venlo, Netherlands). Amplicon length and concentration was analysed using HS DNA 1000 with TapeStation 2200 (Agilent) and Quanti-IT (ThermoFisher) with ClarioStar (BMG). Region specific amplicons were individually barcoded using Illumina’s Nextera XT Index V2 kit with limit cycle PCR and PE150 sequenced on Illumina’s Miseq. The reads were mapped using the ViralProfiler-Workflow, an extension of the CLC Genomics Workbench (Qiagen, Germany). Consensus sequences were generated by a reference-based method and minority variants were called using a cut-off of 1%. Minimal coverage at position PB2-627K was 2000 reads.

### Statistics

For statistical analysis of the virus replication curves, we considered two levels of dependency in the generated data: repeated measures nested within a replicate and replicate nested within a virus-cell combination. To account for these dependencies we fitted linear mixed models (LMM), where this nested structure and repeated measures were included as random effects. The virus titer (TCID50/ml) was the response variable, the variables Time (time of harvest post infection), Virus (two isolates PB2-627E and PB2-627K), Cell (MDCK, A549 or DF-1) and Temperature (37°C or 33°C) as well as their interactions were assessed to evaluate their significance. If an interaction was not significant, it was excluded from the final model. Significant interactions (p < 0.05) kept in the final model were Time:Virus, Time:Temperature and Time:Cell. Variable significance was assessed using the Likelihood Ratio test. This analysis was done using the statistical software R (30), the LMM was fitted using the package lmer (31), and post hoc test for pairwise comparisons were done using the package emmeans (32). Further assessments were done for the data generated from the cell line DF-1. Because of the limited number of observations, this analysis was done by fitting linear models assuming observations were independent (see Results).

### Accession numbers of viruses

The virus sequences generated in this study were submitted to the GISAID database, and accession numbers are listed in Table S2.

## ACKNOWLEDGEMENTS

We acknowledge Albert G de Boer, Arno-Jan Feddema, Corry H Dolstra, Eline Verheij and Frank Harders for technical assistance. We acknowledge Latoya Siemons VRC Zundert, St. Dierenambulance Kennemerland, Dierenkliniek Castricum, Faunabeheer Middachten, and the Netherlands Food and Consumer Product Safety Authority (NVWA) for notifying and submitting the foxes. We thank Wim H M van der Poel for critical reading of the manuscript. This research was funded by the Dutch Ministry of Agriculture, Nature and Food Quality (project WOT-01-003-096 and KB-37-003-039).

## SUPPLEMENTALS

**Table S1: Immunohistochemistry and histopathology**.

**Table S2: Geographical and nucleotide distance between foxes and wild birds infected with HPAI**.

**Figure S1:** Phylogenetic tree for all eight virus RNA segments obtained with the Maximum Likelihood method showing the viruses detected in the samples of the three foxes (orange), and closely related viral RNA sequences from other viruses detected in the Netherlands (green), Europe (blue) and relevant sequences form the 2020-2021 epizootic (blue).

**Table S3: MLL analysis of variables time and virus variants 627E-627K**.

**Table S4: MLL analysis of variables time and temperatures 33°C and 37°C**.

**Table S5: GISAID accession numbers**.

**Table S6: Primer sequences with Illumina tag for amplification of PB2 protein region**.

